# Proximity as a Ground-Truth Proxy for Training Texture Discrimination and Segmentation

**DOI:** 10.64898/2026.05.12.724620

**Authors:** Wilson S. Geisler

## Abstract

Perceptual systems in humans and many other animals are able to segment scenes into regions that are likely to be physically meaningful. This ability depends on having low-level mechanisms that can accurately categorize whether local image patches are samples from the same or different kinds of texture. We find that using spatial proximity as a proxy for same-different ground truth makes it possible to train accurate decision variables and bounds directly from arbitrary natural images with no feedback. We also find that performance can be further improved by using proximity as a ground truth for adjusting the final decision variables and bounds for the current image/scene. These surprising findings result from the simple fact that under a wide range of conditions proximity discrimination (near vs. far) and texture discrimination (same vs. different) have mathematically identical decision bounds if the same image features are used for both tasks. We used the decision variables and bounds trained on natural images as the initial steps in a hierarchical Bayesian observer (HBO) model of texture discrimination [9]. Given the relative simplicity of this HBO model, it did an excellent job of segmenting images having randomly shaped regions containing arbitrary natural textures. We suggest that the proximity proxy is something that natural selection could discover and exploit for any same-different task where the task-relevant stimulus features also vary systematically with distance in space and/or time. For example, natural selection could have created developmental learning/plasticity mechanisms that exploit the proximity proxy.

## Introduction

Many perceptual tasks involve the sub-task of estimating how likely local sets of features are of belonging to the same category versus different categories. For example, segmenting an image into regions likely to contain the same texture or material depends strongly on local same-different categorization, especially if the textures, and the shapes of the regions they define, are unfamiliar. Similarly, local same-different categorization is required for segmenting images into distinct regions of illumination, motion and depth, and for segmenting sound waveforms into sources.

The mechanisms of segmentation in the human perceptual systems are a sophisticated combination of low-level mechanisms that do not depend on prior learning as well as higher-level mechanisms that depend on the recognition of objects, materials, sound sources and scene context. The low-level mechanisms are crucial because they are necessary for training and activating the recognition mechanisms.

Our recent efforts have been directed at characterizing and understanding the low-level mechanisms of texture segmentation [9]. To do this we first defined a family of images we call grown-texture-region (GTR) images that are created by growing randomly shaped texture regions and then filling those regions with randomly selected textures. Figure 1 shows examples of GTR images filled with Brodatz [2], Fabric [9], VisTex [15], McGill [16] and PerTex [4] textures (see Supplementary Material for thumbnails of the texture sheets used from these datasets.)

**Figure 1.**
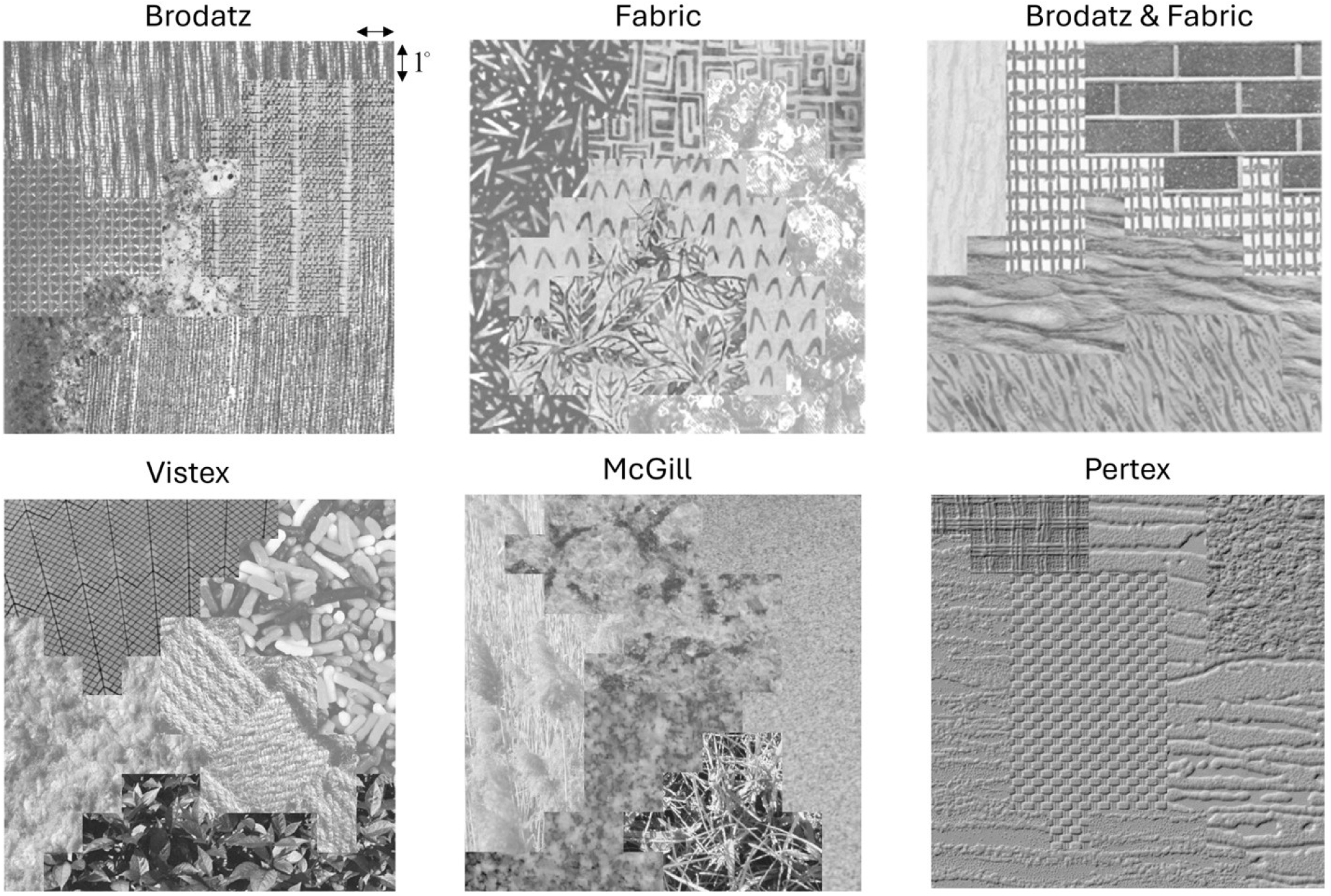
Examples of grown texture region (GTR) images. The regions are grown using random Markov processes and are filled with randomly selected textures [8]. Here the textures are from the Brodatz [1], Fabric [8], VisTex [13], McGill [14] and Pertex [4] databases.

We then defined a biologically plausible set of general and task-specific features and derived, using natural scene statistics and efficient coding, a Bayesian observer model for categorizing pairs of 1° × 1° image patches as being from the same or different texture sheets. This patch size was picked to represent the maximum size of receptive fields in primary visual cortex near the center of the visual field, which is a plausible scale for local image analysis.

We first trained the model to perform as accurately as possible on pairs of texture patches sampled from Brodatz and Fabric textures. We found that this Bayesian observer accurately predicted human ability to categorize pairs of Brodatz texture patches as same or different, as a function of retinal eccentricity. Next, we combined this model of local texture categorization with several fundamental principles of perceptual grouping to obtain a biologically plausible model of texture segmentation. This hierarchical Bayesian observer (HBO) model did an excellent job of segmenting GTR images having regions filled with either Brodatz or Fabric textures, suggesting that the local features, grouping principles, and decision rules applied quite generally. However, when we applied the discrimination model to certain other kinds of texture, such as the Pertex textures (see Figure 1), we found that it performed poorer in same-different patch categorization, leading to much poorer segmentation. This was not due to poor separation of feature responses to the same and different patches, but to the final step of the decision process being nonoptimal. This raises the interesting hypothesis that the human visual system is doing something more sophisticated for learning and adjusting its decision rules.

Here we show that it is possible to accurately learn, directly from natural images, the general rules for same-different texture discrimination by using spatial proximity as the ground-truth reference for learning the decision rules. This is an extension of an earlier less successful approach we tried with color and proximity [6]. We further show here that the final decision bound used for each GTR image can also be appropriately adjusted by using proximity as ground truth.

We find that with the adjusted bounds the segmentation of GTR images becomes optimized. We conclude that perceptual systems may have evolved general default segmentation mechanisms, as well as mechanisms for dynamically adjusting the decision bound for each image/scene, by exploiting the proximity information directly available in the retinal and cortical representations. We also argue that this strategy should work for any task where the task-relevant features vary robustly with proximity in space and/or time.

## Results

### Fundamental principle

The fundamental principle underlying the use of proximity as a proxy for ground truth is surprisingly simple. Consider the GRT image in Figure 2A. Nearby patches are most often the same kind of texture but sometimes they are from different textures. Conversely distant patches are most often from different textures but sometimes they are from the same texture. Thus, the task of judging whether patches are the same or different textures is correlated with the task of judging whether the patches are near or far from each other. The surprising fact is that under many circumstances the decision rules that optimize categorization of proximity (near vs. far) are the same decision rules that optimize same-different categorization.

**Figure 2.**
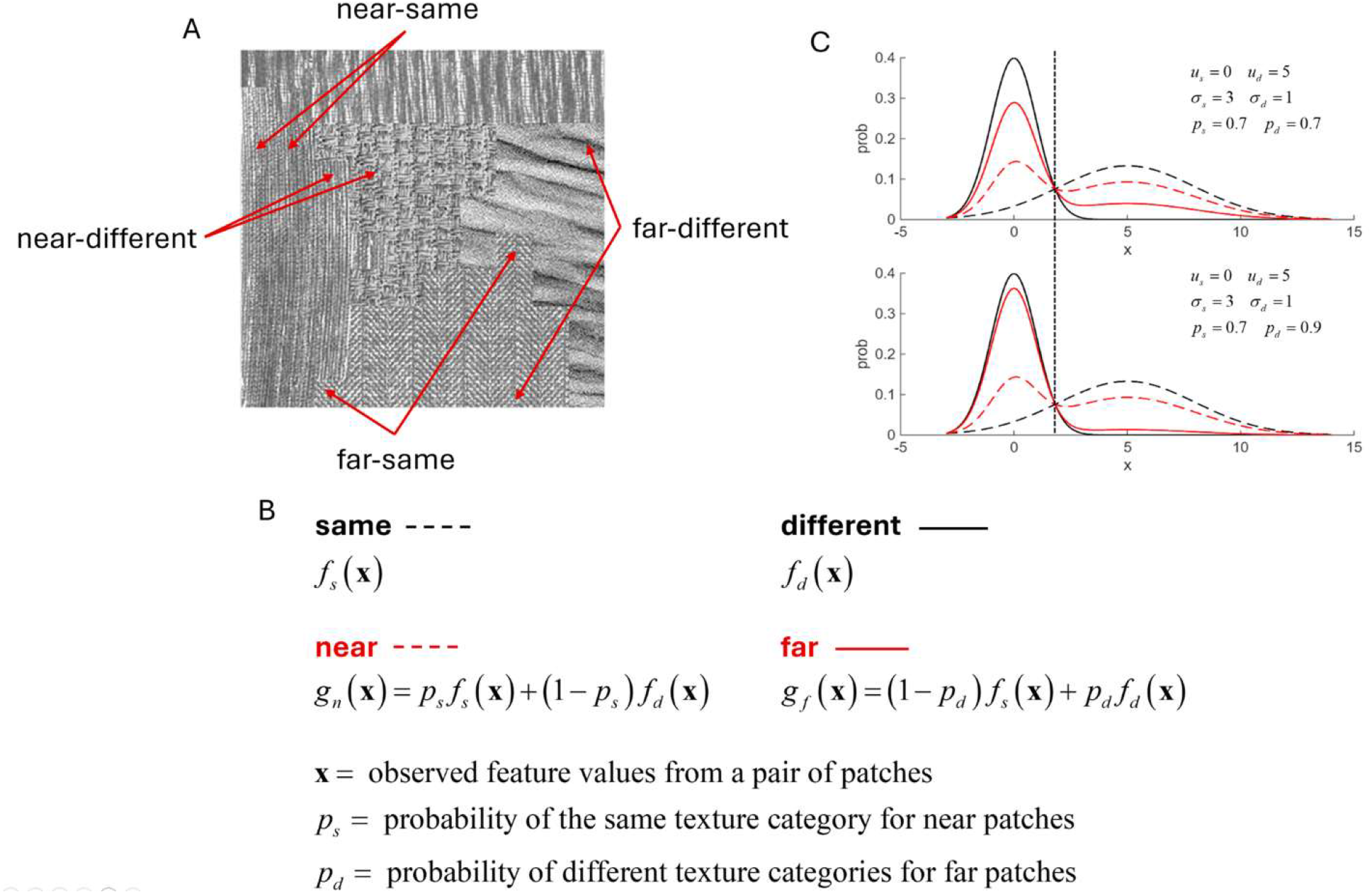
Rationale for using proximity to train for same-different texture categorization. **A.**Both nearby and distant texture patches may contain the same or different texture textures. **B**. The probability distributions of features of nearby and distant texture patches are a mixture of the probability distributions of features from same and different textures. C. Illustration that the optimal decision bound for near-far categorization is the same as for same-different categorization. Parameters are the same-different means and standard deviations, and the mixture parameters.

To see this, let **x** be the vector of task-relevant feature responses from a pair of texture patches in an image, and let *f*_*s*_ (**x**) and *f*_*d*_ (**x**) be the probability distributions of those feature vectors for the same and different categories (see Figure 2B). When the two patches are next to each other there is a relatively high probability *p*_*s*_ that they are from the same category and a probability of 1 – *p*_*s*_ that they are from different categories. In other words, the probability distribution for near feature vectors is a mixture of the same and different probability distributions. Similarly, when the two patches are far from each other there is a relatively high probability *p*_*d*_ that they are from different categories and a probability of 1 – *p*_*d*_ that they are from the same category. Thus, the probability distribution for far feature vectors is another mixture of the same and different probability distributions.

When the prior probabilities of the two categories are equal (or are made equal for training), the decision bounds that optimize accuracy are located where the two probability distributions have the same value. Setting the near and far probability distributions equal to each other, we have (*p*_*s*_ + *p*_*d*_ –1) *f*_*s*_ (**x**) = (*p*_*s*_ + *p*_*d*_ –1) *f*_*d*_ (**x**) and hence *f*_*s*_ (**x**) = *f*_*d*_ (**x**).

In other words, the location of the near-far (proximity) decision bound is identical to the location of the same-different decision bound. Thus, if these assumptions about the relationship between same-different categories and near-far categories are approximately correct, then finding the decision variable and bound that maximizes near-far accuracy will also maximize same-different accuracy. We note that this result is fairly general in that the same-different probability distributions are arbitrary, the feature vectors are arbitrary, and values of the mixture parameters *p*_*s*_ and *p*_*d*_ are arbitrary. If the training is done on a per image basis the mixture parameters can vary arbitrarily from one image to the next.

The fundamental principle is illustrated in Figure 2C for mixtures of one-dimensional Gaussian distributions with different means, standard deviations and mixture parameters. The black curves show the distributions for the same-different categories and the red curves for the near-far categories. The dotted vertical line shows the optimal bound that is between the two distributions. (There is another bound not shown off to the left because the variances for same and different are unequal.)

### Application to same-different categorization of texture patches

If the assumptions in Figure 2 hold for natural images, then it should be possible to learn the general decision rules for same-different categorization, given features **x**, by training on near-far categorization. To test this hypothesis, we trained near-far categorization using a collection of 396 high-resolution calibrated natural images (4,284 × 2,844 pixels) that are 14 bits per color and that were converted to grayscale. The images included natural outdoor scenes, urban outdoor scenes, and indoor scenes. (The images and camera calibration procedure are available at natural-scenes.cps.utexas.edu.) A region from one of the natural outdoor scenes is shown in Figure. 3A.

**Figure 3.**
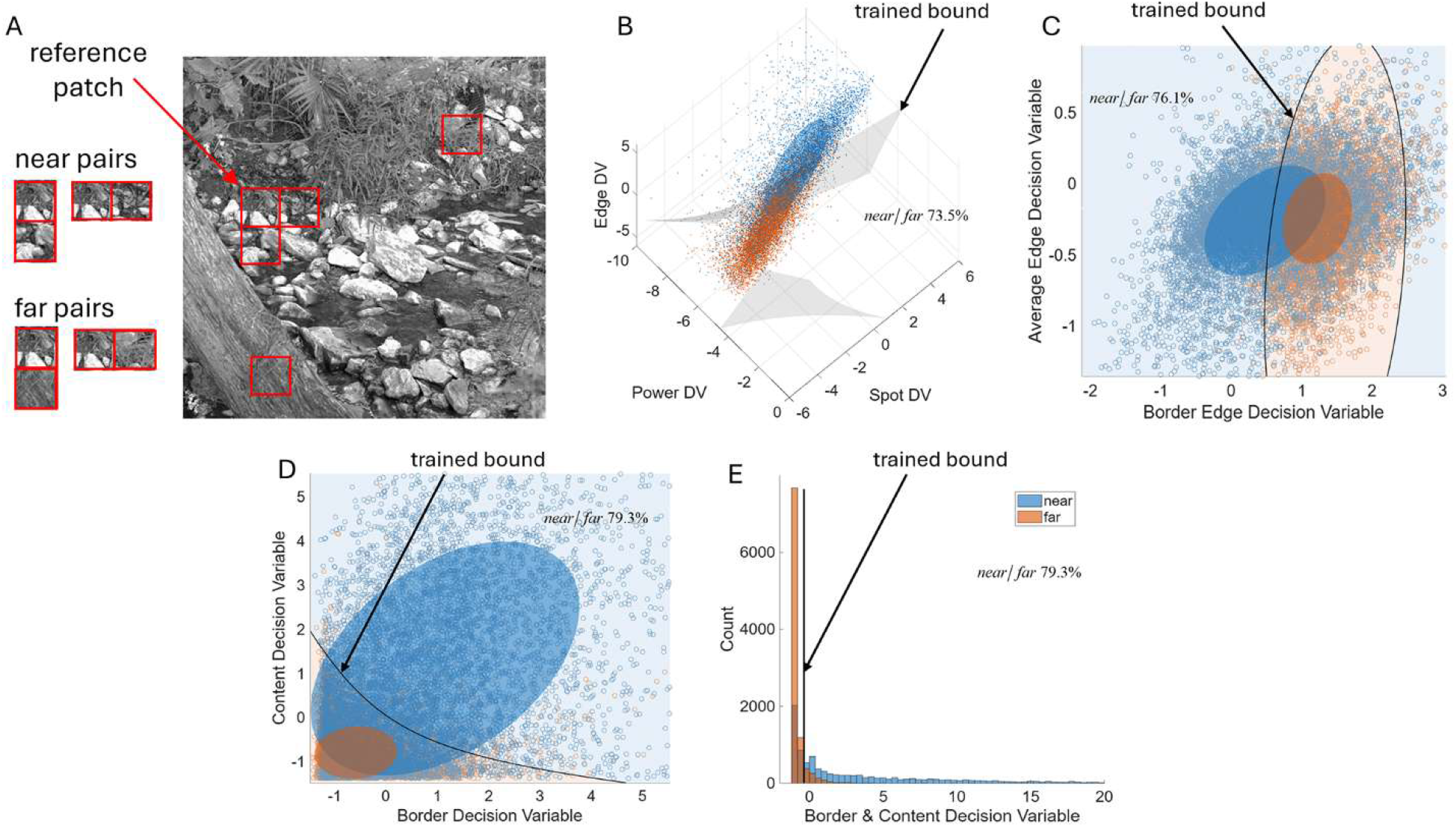
Training decision variables (DVs) and bounds for texture discrimination directly from arbitrary natural images using proximity as ground truth. **A.** Example of near and far pairs of patches sampled from a gray-scale natural image. **B**. Near-far decision bound (for the “content” DV) learned given the values of the near-far decision DVs for three different kinds of task-relevant features: power (complex cells), spot (center surround simple cells), and edge (edge and bar selective simple cells). **C.** Near-far decision bound (for the “border” DV) given the near-far DVs for two kinds of task-relevant feature. **D**. Near-far decision bound given the near-far DVs associated with the learned content and border decision bounds shown in B and C. **E**. The final decision bound can also be represented by a single criterion on the combined border and content decision variable axis.

To simplify the presentation, we consider here a fixed resolution visual system having the same optics and spatial sampling as found near the line of sight in the normal human visual system. However, the proximity proxy for ground truth can also be applied in variable resolution (foveated) visual systems (see Supplementary Material).

After applying a filter matching the optics of the human eye [19], we randomly selected from each natural image ten reference patches that were each 1° × 1° (64 × 64 pixels). The lower and right neighboring patches were used to form two near pairs. Two randomly selected patches beyond a criterion distance (11o) were combined with the reference patch to form two far pairs. Thus, the training was on a total of 7920 near and 7920 far pairs of patches. Consistent with early visual processing, the patches were luminance and contrast normalized [1,12,3].

The model of same-different categorization is a Bayesian observer model which is summarized in the Supplementary Material and described in detail in [9]. The model is hierarchical. The generic task-independent features at the initial level are like those found in the primary visual cortex of primates: edge features (simple-cell edge and bar receptive field responses), spot features (simple-cell center-surround receptive field responses), and power spectrum features (like complex cell responses). These initial features are combined into task-dependent edge, spot and power decision variables which were trained by maximizing near-far categorization. Each of these decision variables represents the log likelihood ratio that the patch pairs are in the near or far category. These decision variables represent the differences in the feature content within the two patches. Each of these decision variables is represented by one of the axes in Figure 3B. Each blue dot represents the responses of a near pair of patches, and each orange dot the responses to a far pair of patches. The gray sheets show the quadratic support vector machine (QSVM) boundary that maximizes near-far categorization accuracy. This boundary is associated with a decision variable that we call the “content” decision variable.

There is also a border decision variable which combines feature responses created at the border between the two patches. Figure 3C shows the learned near-far decision bound for the border features. This boundary is associated with a decision variable we call the “border” decision variable.

After training these bounds and decision-variables, we trained a final decision variable and decision bound (black curve in Figure 3D) that combines the content and border decision variables. After the combined decision variable is learned, the final decision bound can be represented by a single criterion on the combined border-content decision variable axis (Figure 3E).

The accuracies of the near-far categorizations are shown in Figure 3, and as expected they are modest (in the range of 70%-80% correct). Of course, one cannot say anything about the same-different accuracy, because there is no ground truth for our natural images.

However, the accuracy can be checked by applying the decision variables and bounds learned from natural images to GTR images where the ground truth is known. The light gray bar in the leftmost pair of bars in Figure 4D shows the average ground-truth accuracy for the six kinds of textures in Figure 1. The overall average accuracy is quite good (92.7%).

**Figure 4.**
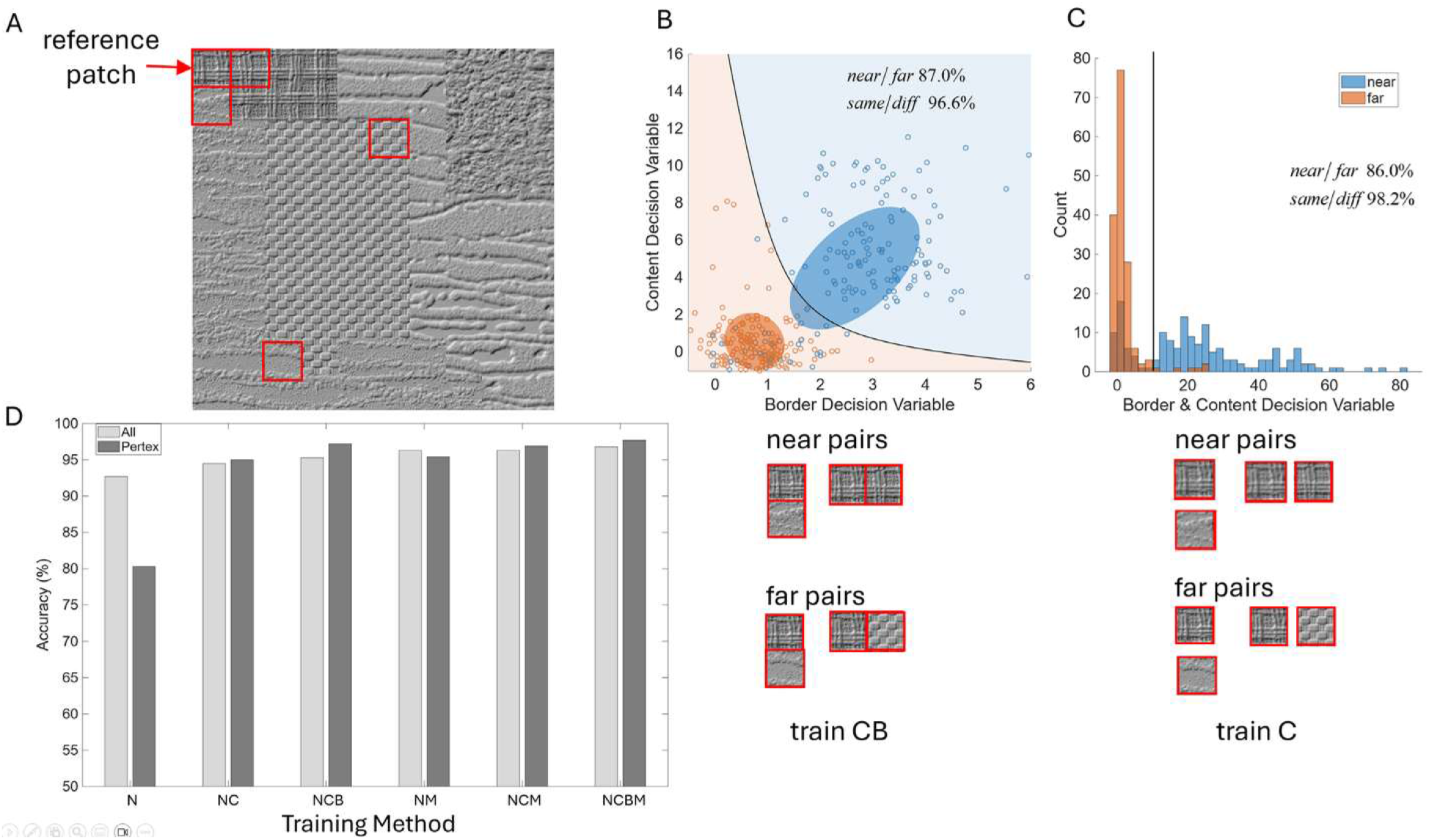
Training the final decision variable and decision criterion for texture categorization from a single image using proximity as ground truth. **A** Sampling near and far patches from a GTR image. **B** Trained decision bound for an example GTR image (NCB). **C.**Trained offset of the content decision variable (NC). **D**. Same-different texture discrimination accuracy for different methods of training. Light gray bars = all kinds of texture. Dark gray bars = Pertex textures. Training methods: N = all training on natural images as in Fig. 3 with no per image training, NC = additional per-image training of a global offset of the content DV, NCB = additional per-image training of the final decision bound, NM, NCM, NCBM = same as first three but with the addition of a final mutual similarity offset.

As mentioned above, when the decision variables and bounds were trained on Brodatz and Fabric textures we found that same-different performance was substantially lower for some kinds of texture such as the Pertex textures. This turned out to also be true for the decision variables and bounds trained on natural images (as summarized in Figure 3). The leftmost dark-gray bar is for the Pertex textures. The performance is so low (80.3%) that segmentation of the Pertex GTR images is very poor.

Thus, we considered possible ways the visual system could use the ground-truth proximity information available in the retinal and cortical representations to adjust the decision variables and/or bounds for each image. We evaluated the five different methods indicated by the labels on the horizontal axis in Figure 4D. The performance of the different methods is ordered by their overall performance (light gray bars). They all improved performance and they all fixed the issue with the Pertex and certain other kinds texture.

Consider first the NCB method. The low- and mid-level decision variables are fixed and trained on natural images (N) using the proximity ground truth as described above. The per image adjustment (CB) is of the final content and border decision bound and criterion (see Figure 3D).

The method of sampling patches for training from a single GTR image is illustrated in Figure 4A. Each GTR image contains 100 1° × 1° patches. The first nine patches in the upper nine rows each serve as a reference patch. The upper left patch in Figure 4A is the first reference patch. For each reference patch, we obtain two “near” patch pairs and two “far” patch pairs. The near pairs are the reference patch and the patch to the left, and the reference patch and the patch below (see pairs in Fig. 4B). The far pairs are the reference patch combined with two patches randomly selected beyond some criterion distance, which is 7° of visual angle. Thus, for each image there is a total of 162 near pairs and 162 far pairs.

In the NCB method we retrained the whole final decision variable and bound for each GTR image (Figure 4B). As in the general training on natural images, the far pairs are still placed side-by-side for training; in other words, the far patches are selected simply to increase the probability that they are from a different texture. The black curve in the upper panel in Figure 4B shows the decision bound learned from a single GTR image. Again, blue dots show the content and border response for near pairs of patches and the orange dots for far pairs. For this example, near-far accuracy is 87% correct and the same-different accuracy is 96.6% correct. The light gray bar in Figure 4D shows the average same-different accuracy for the six different kinds of GTR images shown in Figure 1. As can be seen, the accuracy is quite high, 95.3% correct. This accuracy is comparable to human performance in the fovea for Brodatz textures and is as good as those obtained by training directly on Brodatz textures using the actual same-different ground truth [9]. The accuracy on the Pertex textures jumps to be higher than the average.

The NCB method works well and could be used in practical applications but there are two aspects of it that lower biological plausibility. One is that learning a two-dimensional decision bound rapidly from a single image and then applying it across the whole image seems a bit complicated for neural circuits. Second, training the whole bound requires computing what the border responses would be for the far patches. This assumption is fine for discovering the fixed general decision rules for natural scenes because they are learned via evolution and learning over the life span. However, actually doing this on the fly in natural scenes seems unlikely.

The simpler, more biologically plausible NC method is shown in Figure 4C. In this method, we assume that all the decision variables learned from natural images are fixed and that the visual system can compute for each image the content decision variable at a distance but not the border decision variable. Thus, what is learned is a single, global additive offset to the content decision variable, which can also be thought of as a global normalization (i.e., a gain scalar on a likelihood ratio is an additive offset of the log likelihood ratio). The normalized content decision variable is then passed to the final fixed decision variable computation. This additive offset of the content decision variable is similar a criterion shift on the final decision axis (Figure 4C). The overall average accuracy with this simpler method is still quite good (94.5%) and accuracy on the Pertex textures jumps to be higher than the average.

All the other methods that were considered include a computation called mutual similarity [9]. Specifically, the content similarity of patches is computed across all of the image patches in some large neighborhood to create a vector of content similarities for each of the two patches to the other patches in the neighborhood. The mutual similarity is the cosine similarity of these two vectors. The idea is that if two patches are similar in content to the same set of other patches in the neighborhood, then they are more likely to be from the same kind of texture. Mutual similarity is a fairly complicated computation, but it may be one that could be implemented efficiently with recurrent lateral or feedback networks. Mutual similarity is computed for every GTR image and is added, with a fixed scalar constant, to the combined border and content similarity. Thus, like content decision variable normalization, it has some biological plausibility and alone can solve the issue with the Pertex textures and certain other kinds of texture (see bars labelled NM in Figure 4D).

Figure 5A shows the performance of the biologically possible NCM method and the less plausible NCBM method for all the kinds of GTR image in Figure 1. As can be seen, they both perform quite well.

**Figure 5.**
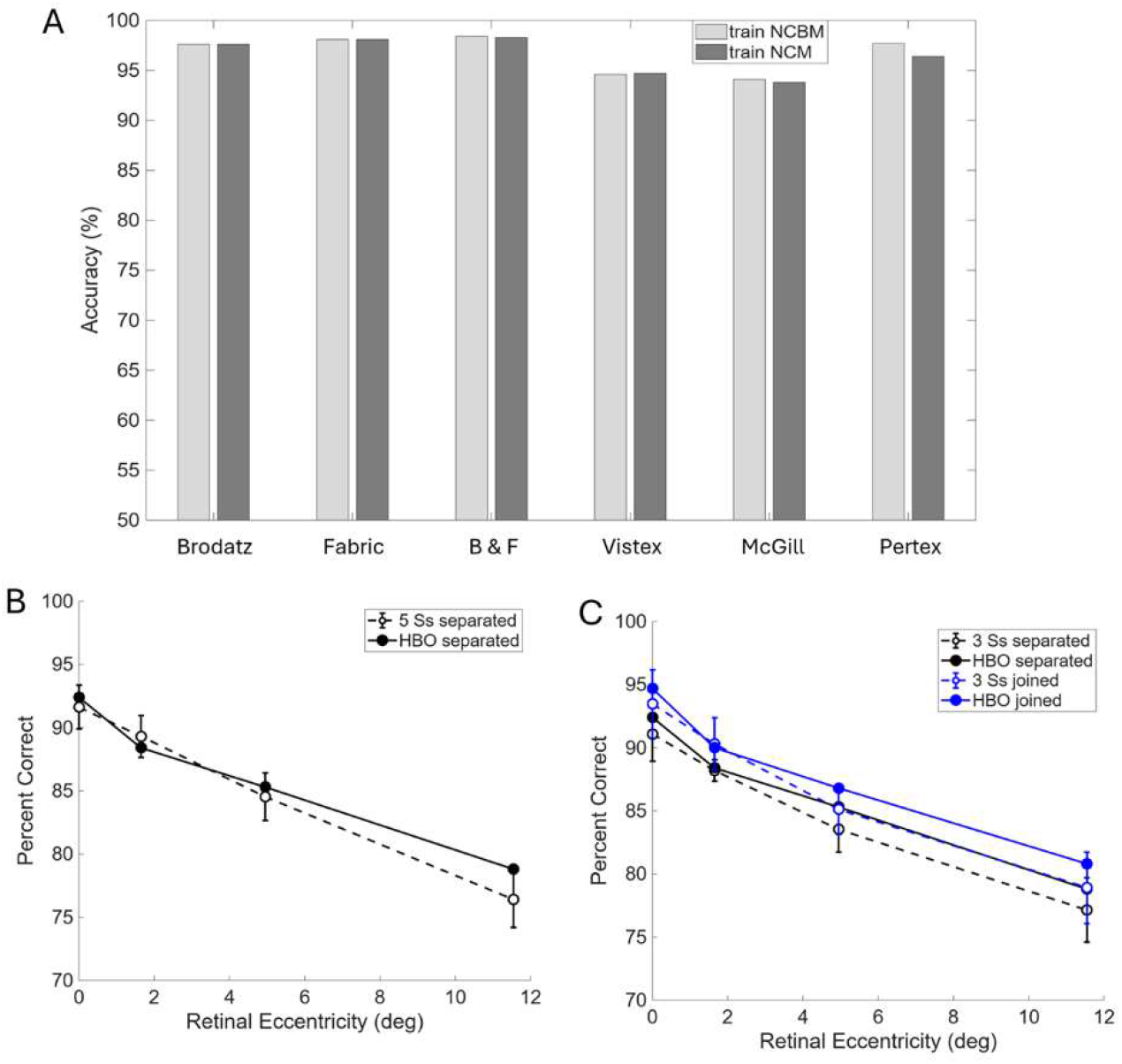
**A**. Performance or GTR images that contain different kinds of texture. **B**. Comparison of discrimination accuracy in joined and separated tasks for human observers and for the HBO model trained by the N method.

As a final note, consider the predicted accuracy in a simple same-different discrimination task for pairs of 1° × 1° Brodatz texture patches presented briefly side by side, as a function of retinal eccentricity [9]. The predictions are based on general decision bounds trained with natural images using proximity as ground truth (Figure 3) separately for each retinal eccentricity, using the N method. The solid black circles in Figures 5B and 5C show the predictions when the two patches are near each other but slightly separated so that only the content information is available. The solid blue circles in Figure 5C show the predictions when the two patches are touching, so that border cues are also available. The open circles in the two figures are the average performance of human observers on these same stimuli [9]. Surprisingly, there is a relatively close match with human performance even though there are effectively no free parameters—the training was only to maximize HBO near-far discrimination accuracy, not to fit the human data. Importantly, these results are almost the same as those we reported earlier when the HBO model was trained on Brodatz and Fabric textures using the actual ground truth [9]. In other words, general training on natural images using proximity as ground truth both performs and predicts human performance as well as training on the specific kind of texture in the experiments while using the actual ground truth. These results strongly suggest that biological vision systems may have evolved by using proximity as a proxy for ground truth in same-different tasks, and that biological vision systems may have also evolved neural learning mechanisms that use proximity as a proxy.

### Application to texture segmentation

The effectiveness and generality of using the proximity proxy for texture discrimination opens the possibility of simple and principled low-level mechanisms for texture/scene segmentation that are effective and general. We tested this hypothesis by using the NCBM and NCM discrimination models as the front end to the HBO model of texture segmentation [9]. The main sequence of computations in this HBO model is shown in Figure 6A.

**Figure 6.**
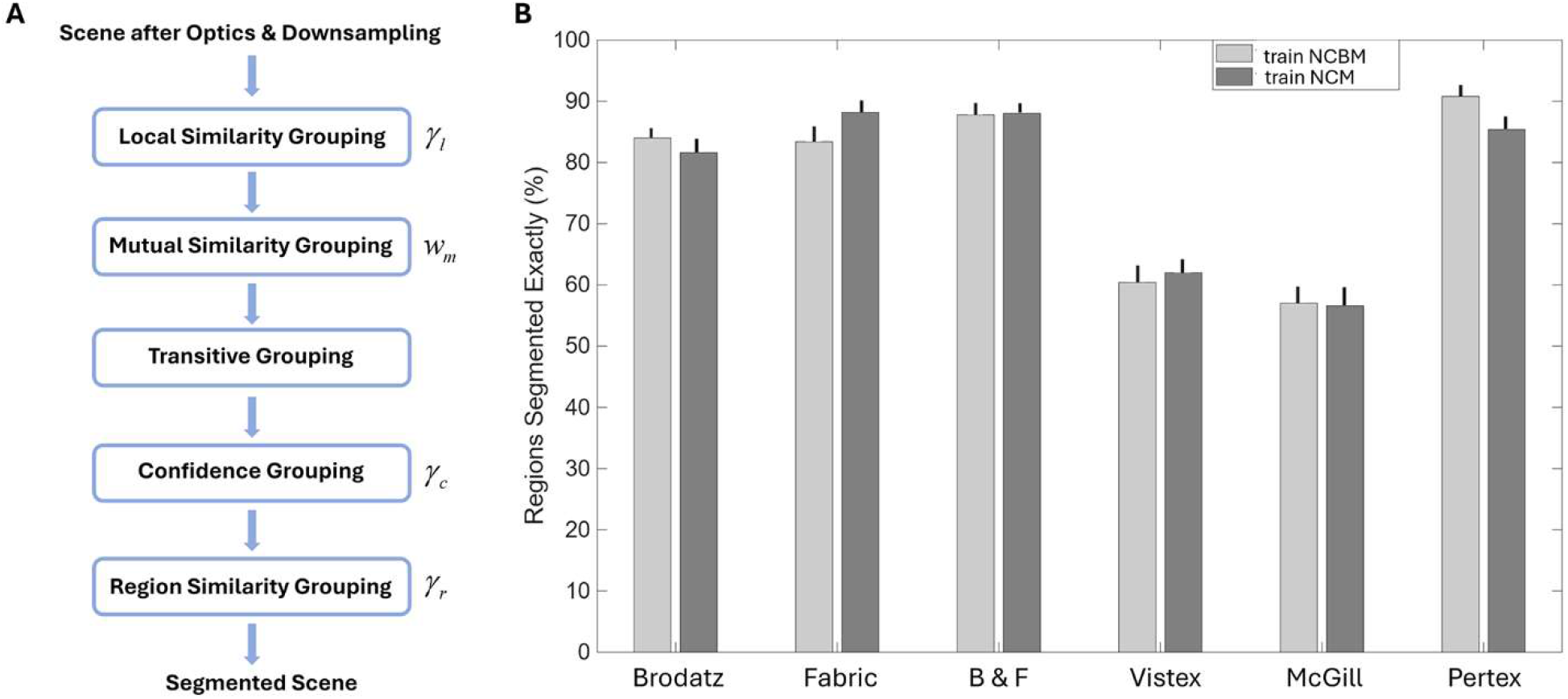
HBO model of texture segmentation. **A.**Main computational steps in the HBO model. There is a single model parameter associated with four of the steps. **B**. Segmentation accuracy (proportion of texture regions segmented exactly correctly) for the six different kinds of GTR images (error bars = standard errors). Parameter values: *γ*_*l*_ = 2.4, *w*_*m*_ = 2.8, *γ*_*c*_ = 0.6, *γ*_*r*_ = 0.8.

The NCBM or NCM models now provide the first and second steps of the HBO model of texture segmentation. The first step is local similarity grouping which links neighboring patches together if they are sufficiently similar [13,16,18,19]. The second step is mutual similarity grouping which boosts local similarity if the two patches are similar to the same set of other nearby patches [9]. The third step is transitive grouping which corresponds to the Gestalt principle of good continuation (continuity): if patch a is linked to its neighboring patch b and patch b is linked to its neighboring patch c, then patches a and c become linked [7,8,11,16,18,19]. This step produces an initial segmentation of the image. The fourth step is confidence grouping [9], which is not actually a single step. Before the transitive grouping step, a confidence criterion on local similarity is applied, so that if a local similarity is near the linking decision bound (i.e., there is low confidence) it is left unlinked and revisited after the transitive grouping step. The final step is region similarity grouping [9] which compares the content similarity of the current groups and merges them if they have a sufficiently high average similarity. There are four parameters in the HBO model (see Figure 6A): the local similarity grouping criterion, the mutual similarity weight parameter, the confidence grouping criterion, and the region similarity grouping criterion.

We applied the HBO model to the six different kinds of GTR images, 120 images each kind. We estimated the four parameters for each kind of GTR image and found that they are similar and hence used the average parameter values which are given in the caption of Figure 6. The segmentation accuracy for all GTR images, using the average parameter values, are shown in Figure 6B. We defined the accuracy to be the percentage of texture regions that were segmented exactly correctly. Across the different kinds of GTR image, the percentage ranged from 50% to better than 80%. For the Brodatz and Fabric textures this is even better than we obtained in the earlier study [9] with a single decision bound trained exclusively on Brodatz and Fabric GTR images. Figure 7 shows examples of GTR images that were segmented exactly correctly for each of the six kinds of GTR images.

**Figure 7.**
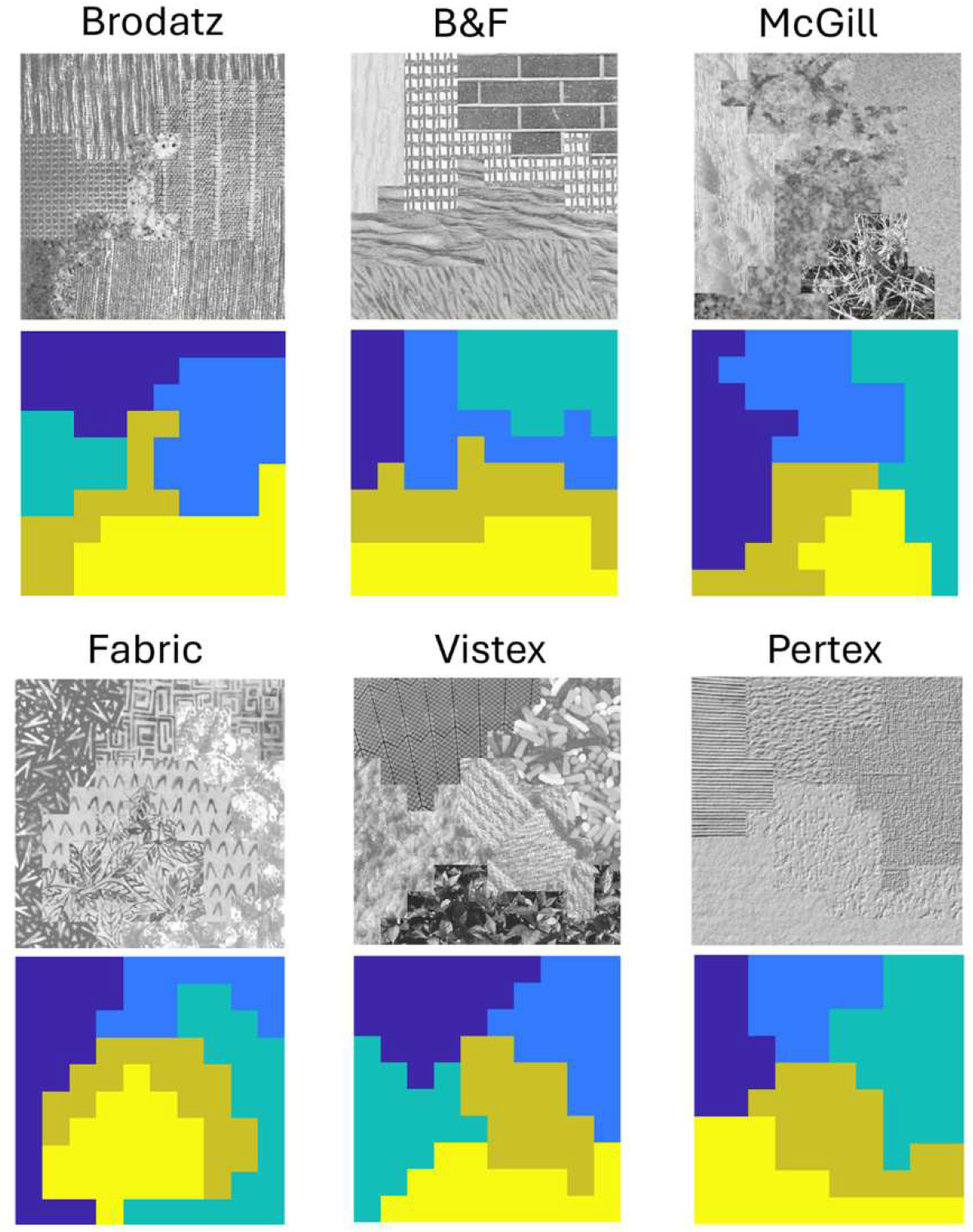
Examples of exactly correctly segmented GTR images. Below each image is an image of the texture regions found by the HBO model, which is also the ground truth.

## Discussion

Perceptual systems in humans and many other animals are able to segment scenes into regions that are likely to be physically meaningful. This ability depends on having low-level mechanisms that can accurately categorize whether local image patches are the same or different kinds of texture. We showed that using spatial proximity as a proxy for same-different ground truth makes it possible to train accurate decision variables and bounds directly from arbitrary natural images. We also showed that overall performance can be further improved by using proximity as a ground truth for adjusting the final decision variables and bounds for the current image/scene.

These findings result from the simple fact that under a wide range of conditions proximity discrimination (near vs. far) and texture discrimination (same vs. different) have mathematically identical decision bounds if the same image features are used for both tasks. This mathematical result follows from the assumption that the probability distributions of the texture features in near and far patches are each a mixture of the probability distributions of the texture features in same and different texture patches.

It is likely that there are other same-different tasks where this assumption holds to good approximation. Patches that are near in the image are likely to have the same illumination spectrum. Thus, decision variables and bounds relevant for same-different discrimination of patch illumination might be accurately learned directly from natural images. Similarly, sound segments that occur near in time are likely to be from the same source. Thus, decision variables and bounds for features relevant for same-different discrimination of sound source, might be accurately learned directly from natural sound recordings.

Proximity in space and time probably played a critical role in the evolution of the general fixed features and computations for same-different discrimination. For example, one likely scenario is that visual systems initially evolved because they could provide some crude information about the direction of light sources (e.g., where is the entrance of a hole in the ground or for aquatic animals which way is the surface of the water?). Proximity then evolved to become a crude cue for grouping locations (directions) into regions likely to be from the same physical source— neighboring locations having similar light levels are likely to be from the same physical source and so are grouped together in neural representations. Then, using proximity as a grouping rule allowed natural selection to discover other features for similarity grouping. The high validity of the proximity ground-truth proxy made this relatively easy because any mutations toward useful spatial and chromatic features had a strong positive effect on local similarity discrimination, grouping, and segmentation.

It is also possible that natural selection found circuits that use the proximity proxy for rapidly adjusting the computations for individual scenes. The present simulations showed that adjusting the gain on the content information decision variable (near-far likelihood decision variable) to maximize near-far discrimination in the image, substantially improved the accuracy and generality of same-different texture discriminations. Adjustments of gain over large neural populations are not uncommon in the brain, e.g., attention-gain changes [3,17], and thus this is a possibility. It is hard to guess the likelihood that such mechanisms exist, but it seems like an important hypothesis to explore. For example, further simulations would be useful to determine the optimal and minimal spatial extent of the computations required for good performance. The minimal and optimal spatial extent would help constrain the possible neural circuits.

The present simulations also showed that mutual similarity, another global computation on each image, substantially improves accuracy and generality. Mutual similarity computations could be implemented with recurrent neural networks and hence it is also possible and hence a useful hypothesis to explore. Again, further simulations to determine the optimal and minimal spatial extent of the computations required for good performance would be useful.

It is also possible that natural selection created slower learning mechanisms that use proximity as a proxy for ground truth. For example, the basic circuits for low-level segmentation may be largely present at birth but may need to be refined and adjusted over time for the infants’ natural environment and other aspects of the developing sensory systems (e.g., development of the eyes’ optics and retina). The spatial and temporal proximity signals, which are easily available in various cortical areas, could serve as the reinforcing ground-truth signal for those refinements and adjustments. Perceptual learning that depends only on the actual ground truth via interacting with the environment would be very slow and sketchy. The proximity proxy would even allow learning from scenes with large distances where motion parallax provides no useful signals.

Finally, our results confirm that combining the principles of Bayesian statistical decision theory with the principles of natural selection is a sensible approach to the study of animal evolution [10].

An interesting and relatively easy direction for future research would be to use the proximity proxy described here to explore the effect of different natural environments, and the natural textures in those environments, on the decision variables and decision bounds for texture segmentation. It would also be interesting to explore what decision variables and decision bounds for texture segmentation to expect for different animal species given their optics, retinal coding, and natural environments.

## Acknowledgements

Supported in part by NIH grants EY11747 and EY024622.

## Supplementary Materials

### HBO model of texture discrimination

Figure S1 outlines the HBO model for texture discrimination, which has four steps.

**Figure S1.**
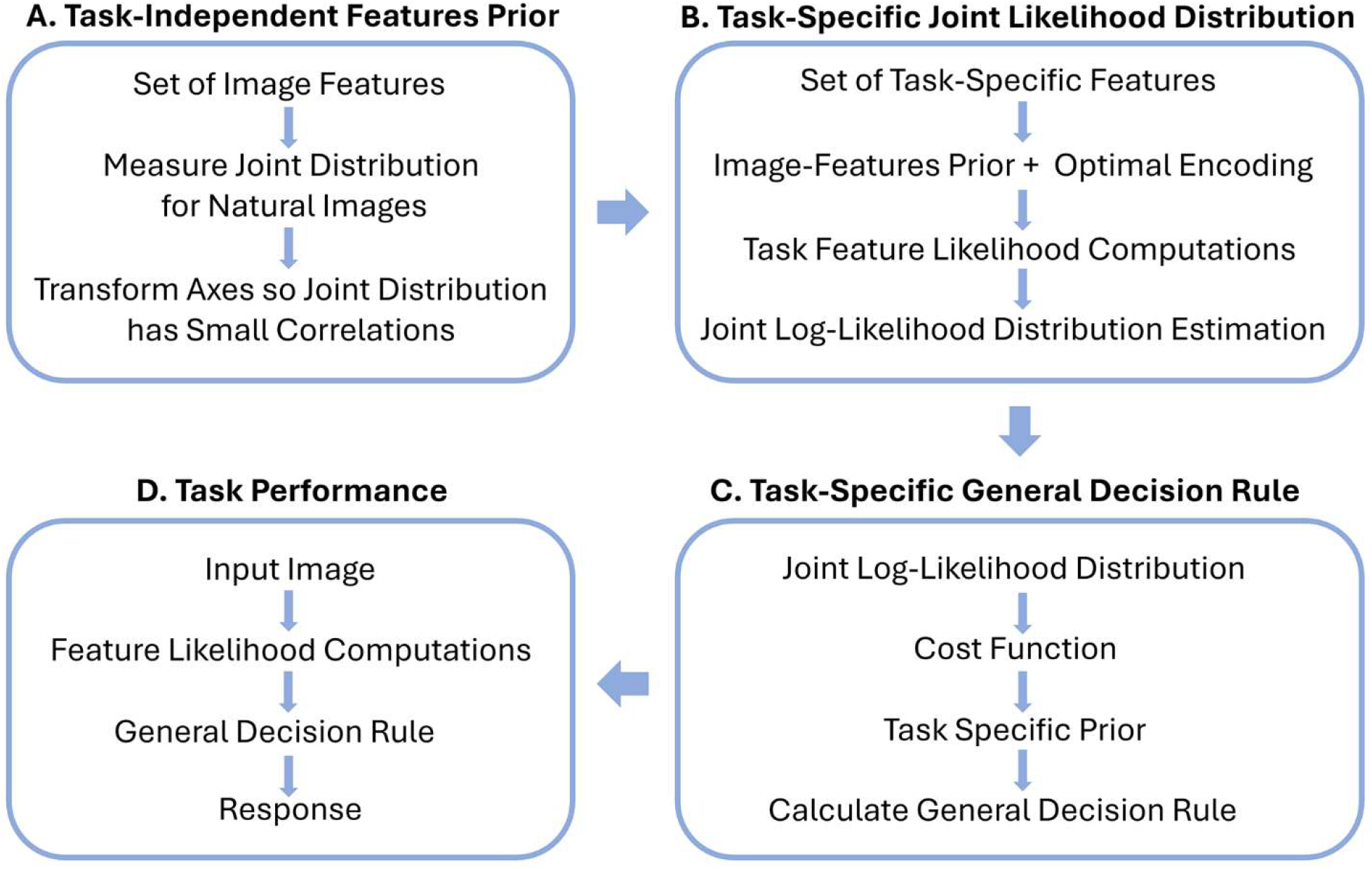
Hierarchical Bayesian Observer (HBO) framework for local similarity grouping (local texture discrimination).

The first step (Fig. S1A) is to specify a set of image features to extract, then determine their joint distribution in natural images, and then, if appropriate, transform the axes so the joint distributions have small correlations. The aim of this first step is to measure the prior distribution of the feature responses largely independent of the specific task. The idea is that evolution and learning over the lifespan will exploit the prior distribution when evolving or learning task-specific features. The task-independent priors do not solve the task, but they are easy to measure, and they provide useful constraints.

The set of task-independent features is show in Figure S2. They consist of (a) steerable 1^st^ and 2^nd^ derivative Gaussian kernels (Figs. S2A and S2B) that are meant to capture the information in the responses of orientation-selective cortical simple cells, (b) non-orientation-selective center-surround kernels at two scales (Fig. S2C) that are meant to capture information in the responses of non-orientation-selective cortical simple cells, and (c) the power spectrum of the 1 × 1 deg patch that is meant to capture the information in the cortical complex cell responses.

Because the texture patches are quite small, the size of the steerable filters were kept as small as possible: 3 × 3 pixels with a standard deviation of 1 pixel. Figure S2D shows the spatial frequency tuning of the two steerable filters and the two center-surround filters. The peak spatial frequency in the fovea is 8 c/deg for the first derivative filter, 11 c/deg for the large center-surround filter and 21 c/deg for the second-derivative and small center-surround filters. These frequencies drop by a factor of two for each factor of two in midget ganglion cell spacing (e.g., peaks become 1, 1.4, and 2.6 c/deg at 11.55° eccentricity).

**Figure S2.**
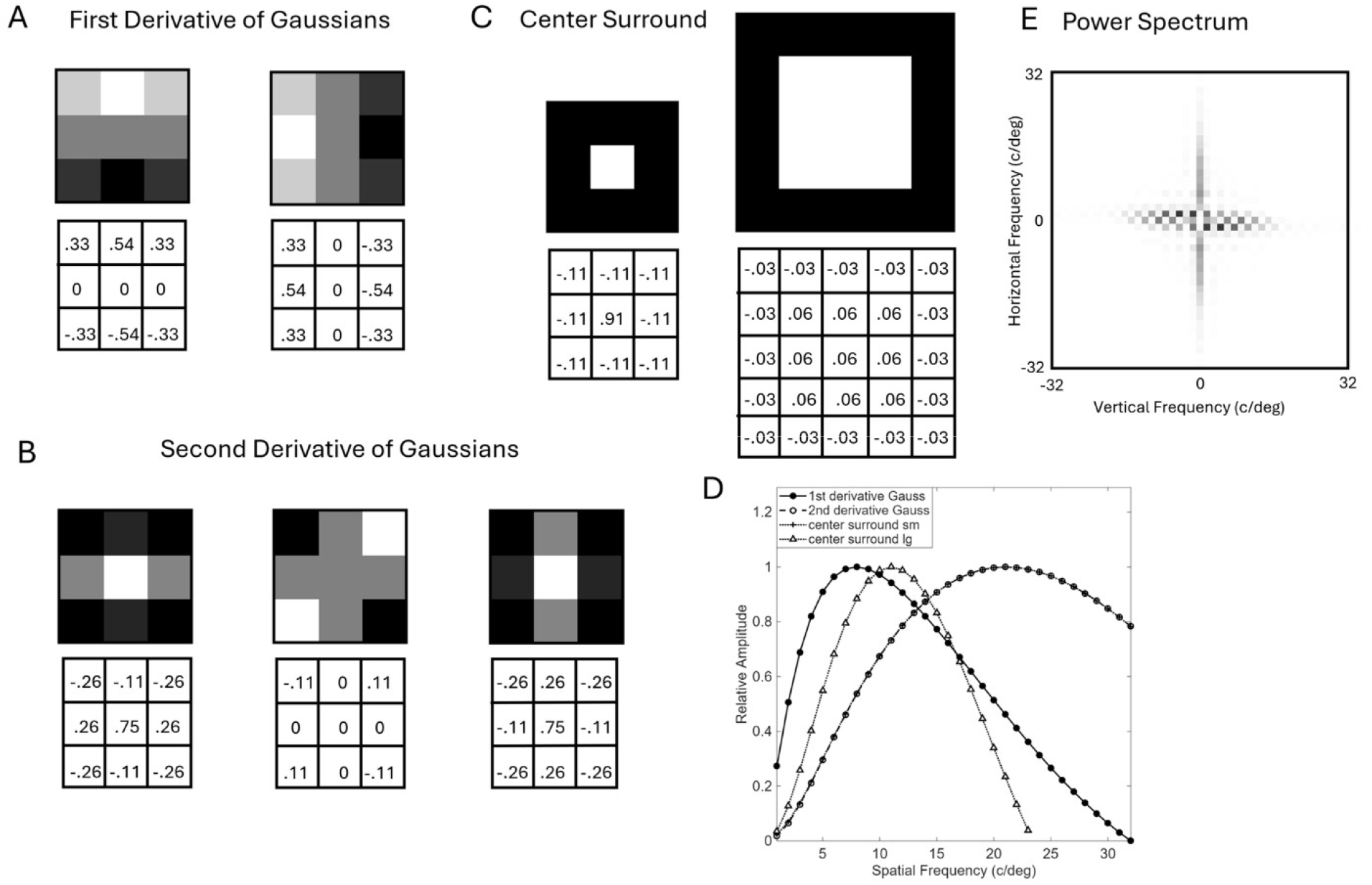
Task-independent features. **A**. Kernels of the first-derivative steerable Gaussian filter. Numbers sre the weights in each pixel **B**. Kernels of the second-derivative Gaussian steerable filter. **C**. Kernals of the center-surround filters. **D**. Spatial frequency tuning curves of the 4 filters in the fovea combined with the effect of the optical transfer function of the human eye. Despite the small kernal sizes of the steerable filters (3 × 3), there is only a small bias in the estimations of the magnitude and orientation of sinewaves. In other words, the steerablity is quite accurate. **E**. Patch power spectrum.

The prior probability distributions of responses of these feature responses were measured for same set of natural images used here for training the decision variables and bounds.

The second step of the HBO model (Fig. S1B) specifies the task-specific features (computed from the input feature responses), and then uses the image-feature prior and principles of optimal encoding when deriving the computations that give the task-specific feature likelihoods and their joint log likelihood distribution. The task-specific features are identical for the same-different and the near-far tasks. Figure S3 lists all the log likelihood decision variables. The individual log likelihoods of same vs. different and for near vs. far for the spot and contour features are given by a histogram decision variable (the first equation). The power spectrum features decision variable is given by the fourth equation.

All the other log likelihood decision variables are the optimal decision variable trained on natural images (or on Brodatz and Fabric textures) using our publicly available software [5].

**Figure S3.**
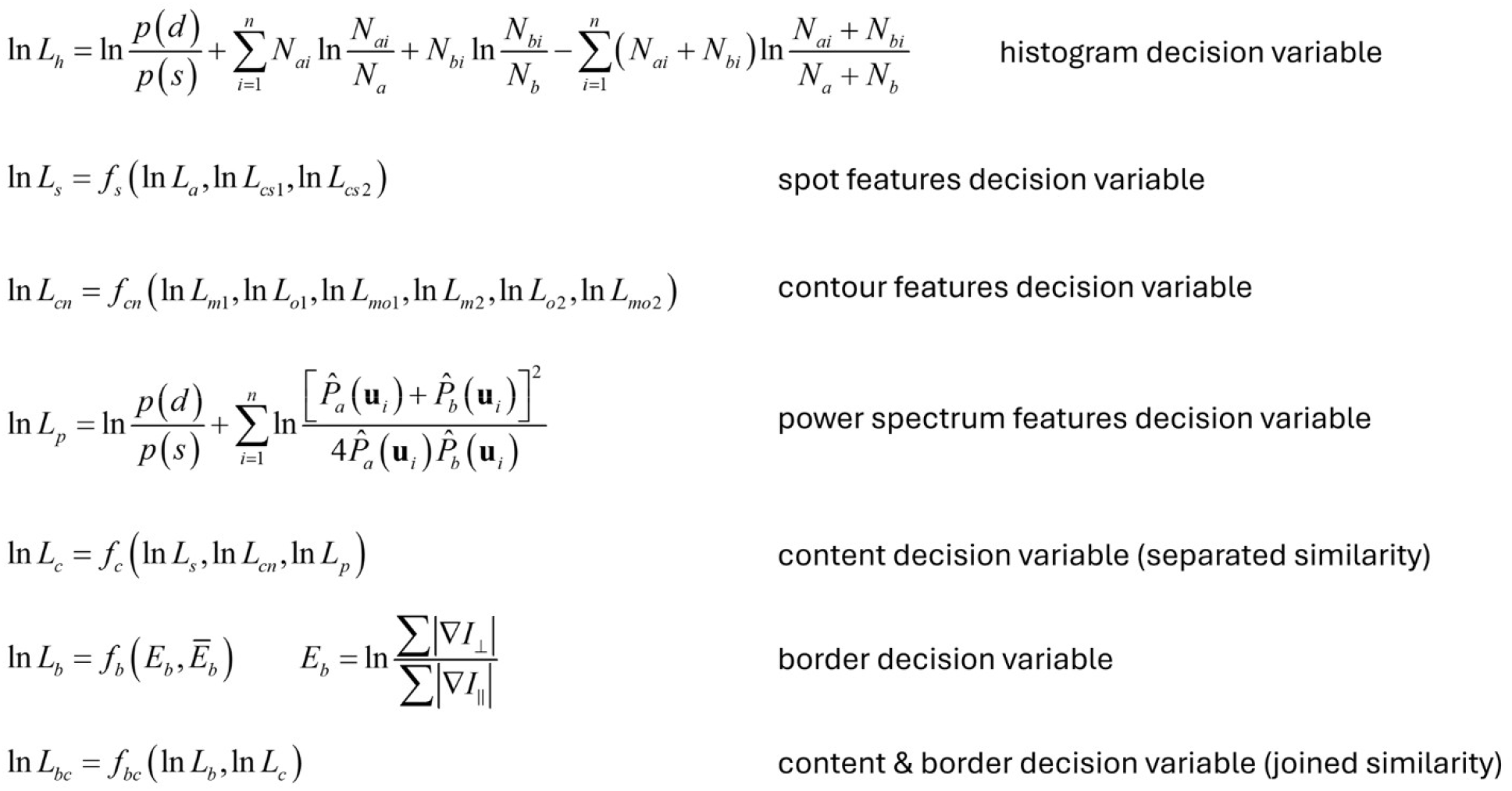
Task-dependent features computed on pairs of patches in the discrimination task and in the image segmentation task. The similarity measure for separated patches is based only on the content features of the patches. The similarity measure for joined patches is based on both the content and border features.

The third step (Fig. 4C) combines the log-likelihood distribution with a cost function and task-specific prior in order to determine the general decision rule. This defines the HBO model of texture discrimination. The fourth step (Fig. 4D) is to simulate the performance of the model. For more details see [9].

All of the above computations and training of the HBO model can be repeated at different retinal eccentricities. A simple computational strategy is to down-sample the input image (e.g., natural image), after filtering with the optics, by successive factors of two which would correspond to the midget ganglion cell spacing [21] at specific retinal eccentricities (e.g., 0°, 1.65°, 4.95°, and 11.55° for the average human visual system). The predictions for other eccentricities can be obtained by interpolation.

### HBO model of texture segmentation

#### Local similarity grouping

The first step of the HBO model of texture segmentation is to apply the above HBO discrimination model to every pair of neighboring texture patches to obtain a same-different (or near-far) log likelihood ratio. The log likelihood ratios can be regarded as similarity measures, the larger the log likelihood ratio the greater the pairwise similarity. Let *φ*_*ij*_ be the log likelihood ratio (similarity) between two patches. When the patches are neighboring, the similarity is based on content and border cues. When they are not neighbors, it is based on only content cues.

#### Mutual similarity grouping

In the next step, the content similarities are computed between all pairs of patches at all distances. The mutual similarity *µ*_*ij*_ between a neighboring pair of patches is defined to be the cosine similarity between the vector of all similarities to patch *i* and the vector of all similarities to patch *j* (the dot product divided by the product of the vector lengths):

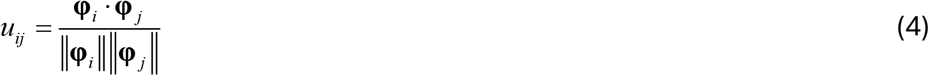

where the bold symbols represent the similarity vectors.

The combined similarity for each neighbor pair is given by

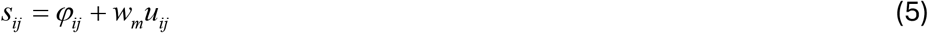

where *w*_*m*_ is a constant that is adjusted to maximize overall performance. Mutual similarity can be thought of as a form of recurrent global feedback to the local similarity between neighboring pairs of patches. Intuitively, if a neighboring pair has the same pattern of similarities to other patches, then they are more likely to be from the same texture region. The combined similarity of each neighboring pair is compared to a grouping criterion *γ*_*l*_. If the similarity exceeds the criterion, the pair is nominally labelled as same, and below the criterion as different. However, similarities that are too close to the criterion are regarded as uncertain. Specifically, if the distance from the criterion is less than some criterion distance, |*s*_*ij*_ – *γ* _*l*_| < *γ* _*c*_, then the link is labelled as uncertain, and not used in the transitive grouping step. This suppression of weak links is a component of confidence grouping.

#### Transitive grouping

The next step applies transitive grouping (a global grouping process) to all pairs labelled “same.” As mentioned in the main text, if patch ***a*** groups with patch ***b***, and patch ***b*** groups with patch ***c***, then patches ***a*** and ***c*** are grouped. Transitive grouping creates an initial segmentation of the image into regions, although there are sometimes a few isolated regions that consist of only one patch. Transitive grouping is a crucial process that allows grouping of textures that are stationary as well as textures that are nonstationary but smoothly varying. In images of the natural environment, smoothly changing textures are common because of perspective geometry and physical processes that grow/create surfaces. Transitive grouping makes segmenting these textures possible. On the other hand, because it causes grouping to propagate it requires some control.

Specifically, an accidentally strong similarity between a pair of patches in two large texture regions can cause the regions to merge into one region (see below).

#### Confidence grouping

Confidence grouping helps to control transitive grouping. Low confidence similarities are places where transitive grouping could inappropriately merge regions together, thus they are ignored in the transitive grouping step. For many low-confidence links, the two patches still end up in regions following transitive grouping, because there are generally four similarities computed for each patch and any one of those could pull the patch into a region. For any patch that is not in any region (a rare event), its content similarity is computed for all patches in the segmented regions that they touch.

They are then assigned to the region with the highest average similarity.

#### Region similarity grouping

This compares average content similarity of the regions obtained with the first four steps. If the average content similarity of two regions exceeds a criterion *γ*_*r*_, then the regions are merged. Occasionally, a region is isolated in that it consists of just one patch. That patch is merged with whatever neighboring region best matches it.

#### Texture Sheet Thumbnails

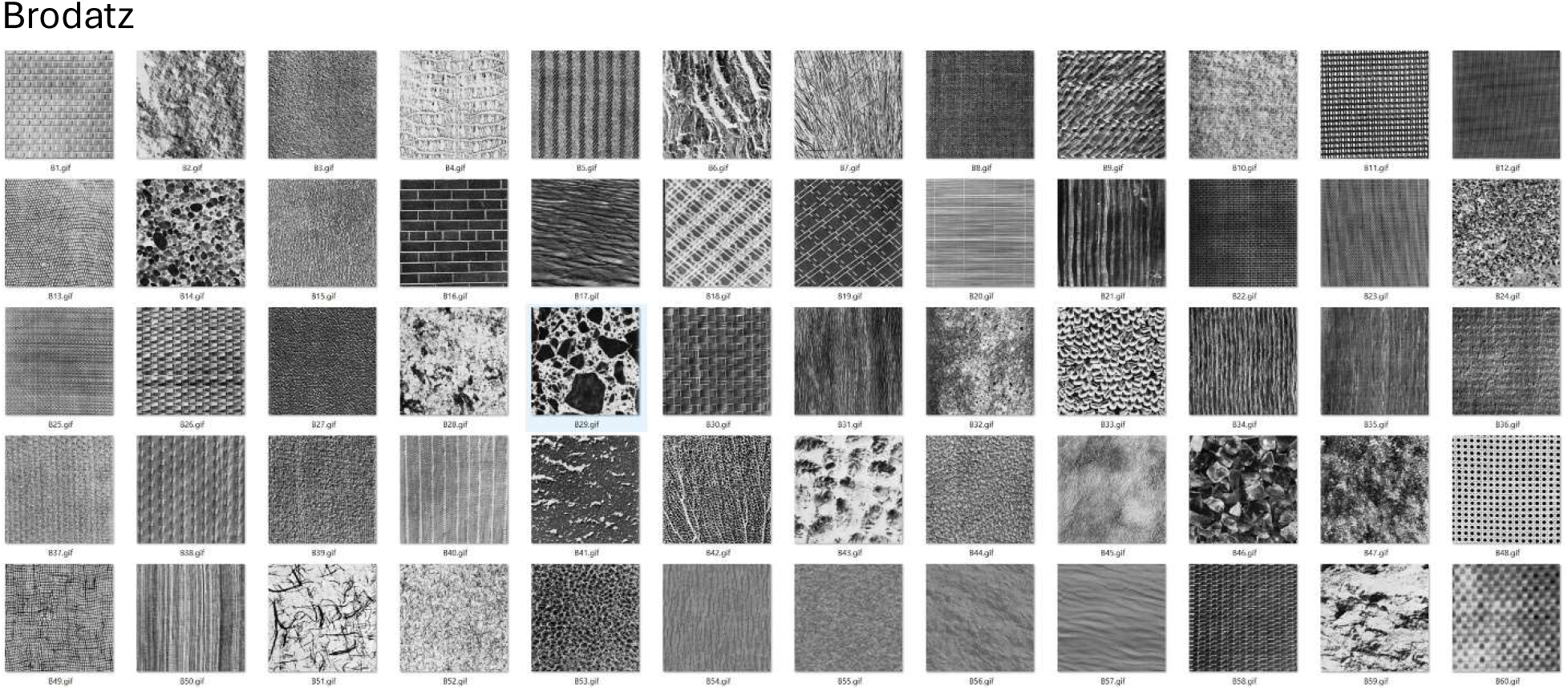
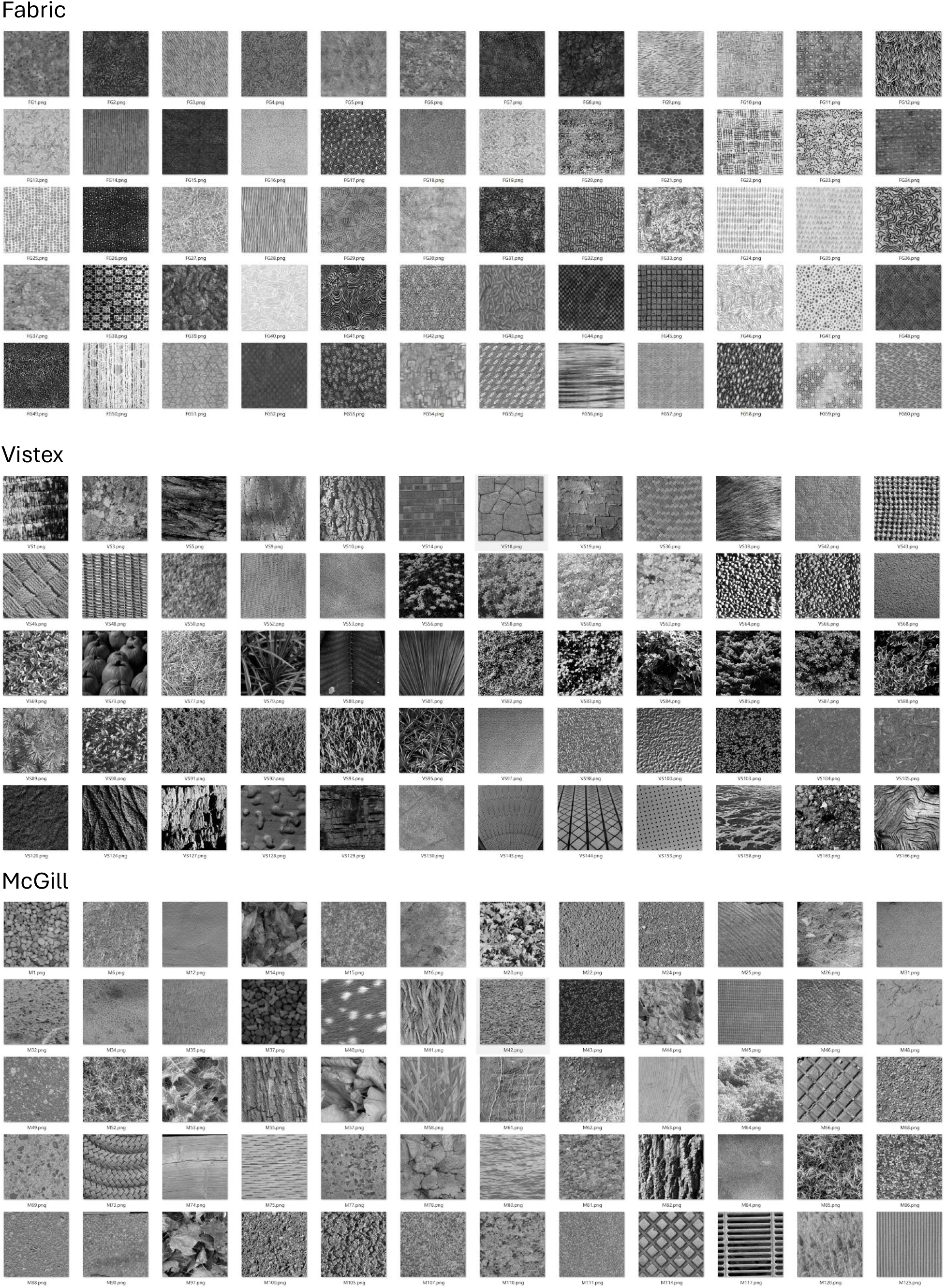
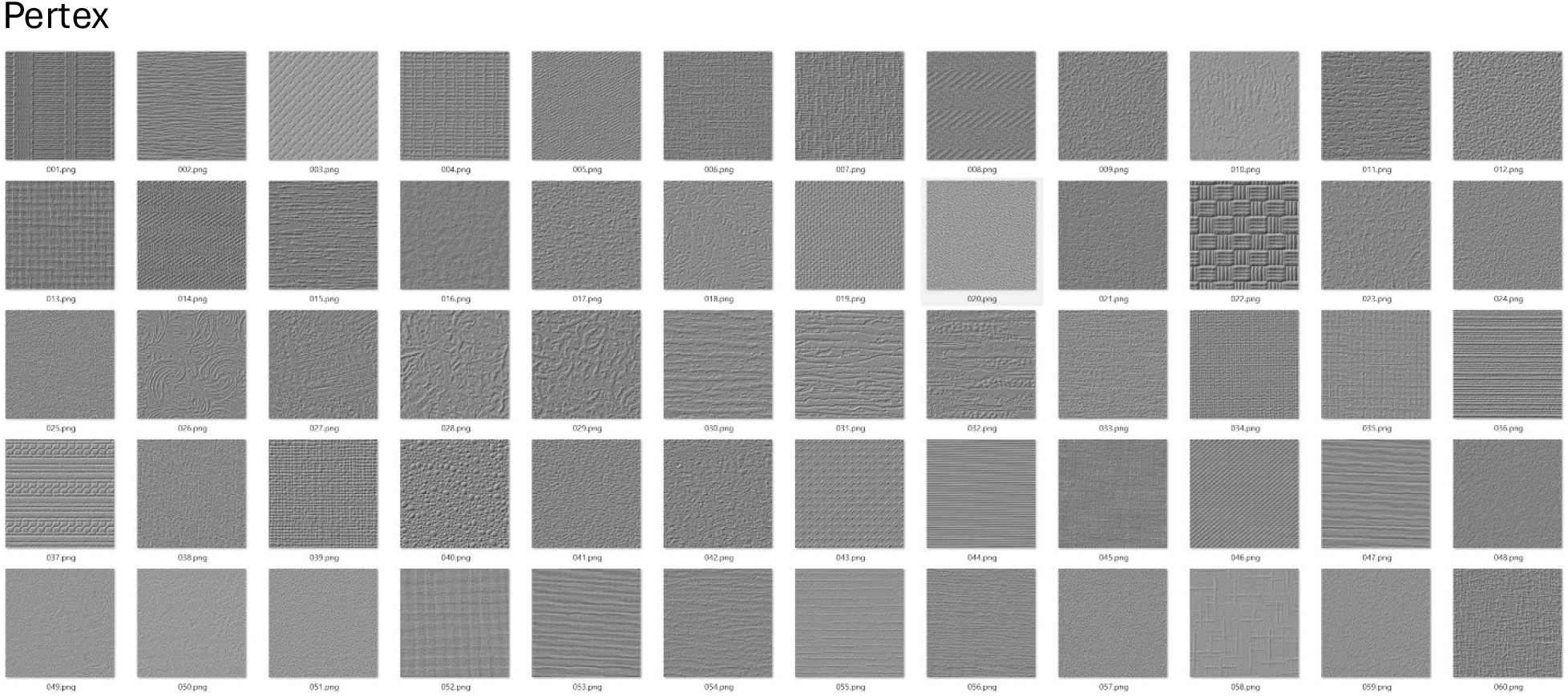

